# An assessment of land use land cover change in Central Highland of Deccan Peninsula and Semi-Arid tracts of India

**DOI:** 10.1101/665794

**Authors:** Tamali Mondal, Parabita Basu, Qamar Qureshi, Yadvendradev Jhala

## Abstract

Human consumption and management of land, i.e., how people transmit to a landscape as a source of livelihood, shelter, recreation, or industry, are powerful forces shaping patterns and dynamics in human-occupied landscapes. Understanding and managing landscape change, therefore, requires a thorough knowledge of the developmental processes the area is undergoing, causing the change in the land use pattern over time. In this study land use/cover dynamics of the Central Highland of Deccan Peninsula and Semi-Arid zone of Gujarat Rajputana bio-geographic regions of India was assessed using a combination of hybrid classification, markov analysis approach within the time frame of 2000-2013. In this time frame, major changes have been identified in dense forest, open forest, cropland, fallow land and in the barren landforms. The landscape has witnessed a substantial increase in built-up areas from 31317 sq. km. in 2000 to 52338 sq. km in 2013. The scrubland has largely declined from 35713.06 sq. km in 2000 to 23003.88 sq.km in 2013; it might have resulted from the transition of scrubland to the fallow land which was predicted as more than 70% in the first phase and more than 30% in the later phase. The area under dense forest constantly decreased in the studied time (84027 sq. km to 71225.37 sq. km). The Markov analysis estimated a probability of changing of open forest types to fallow land (36% in 2000-2006 and 20% in 2006-2013). As the environmental damage is irreversible due to industrialization, therefore a sound strategy needs to be implemented for sustainable use of natural resources that can go hand in hand.

## Introduction

Conversion, degradation and fragmentation threaten the integrity of ecosystems worldwide (Uddin K et al. 2016). Farmland expansion, deforestation and urbanization are the major developmental activities in the Anthropocene epoch causing worldwide depletion of biological diversity at genetic, species and ecosystem levels (Pardini R et al. 2017; Jenkins 2003; Caswell and Cohen 1993; Turner 1987; Miller 1982; Connell 1978). Nowadays, biological species live in increasingly fragmented habitat islands embedded in a matrix of human civilization (Fearnside 2001, Sagan et al. 1979; Charney et al. 1975; Otterman 1974) where the change in land cover is evident in a heterogeneous cluster of ecosystem called a landscape. The landscape, as defined by Forman and Godron (1986); is a heterogeneous land area composed of a cluster of interacting ecosystems that are repeated in similar form throughout. Diverse land use activities like human settlements, agriculture, livestock raising, forest harvesting directly affects the land cover of an area and this is a global phenomenon today (Laurance 2008; Sanderson & Harris, 2000; Xiao et al. 1991). Shifting in the land use patterns caused by social and economic issues; results in land cover change. Other than anthropogenic drivers land cover can be changed by natural events such as weather, flood, fire, climate fluctuations, and ecosystem dynamics (Townsend et al. 2008) which gets further reflected in the landscape structure over time (Castillo et al. 2015; Baker et al. 2015; Otero et al. 2015) in different spatial sizes and frequencies as well (Farina 2000). Apart from the information of existing land cover changes; it is indeed important to monitor environmental dynamics of the land under influence of increasing population which changes the shape and nature of the landscape over time.

The ability to map landforms is an important aspect of any environmental or resource analysis and modelling effort. To ensure a reliable quality of results several important steps need to follow. Conventional ground methods of land use mapping are labour intensive, time-consuming and are done relatively infrequently (Prakash A & Gupta R. P. 2010; Rogan J. 2004). These maps soon become outdated with the passage of time, particularly in a rapidly changing environment. The traditional method of data collection of land use/cover through the survey is quite difficult (Rao D. P.; Gautam N.C.; Nagaraja R & Mohan P. R. 1996; Olorunfemi 1983). Nowadays the developed techniques of satellite imagery processing by remote sensing and GIS analysis have cut down the cost and time to prepare land use/cover maps in regular intervals of time (Mc Garical et al. 2002). Remote sensing and GIS techniques have the capability to record a response which is based on many characteristics of the land surface, including the natural and artificial cover where the structure of a landscape is defined by the spatial pattern and represented by two components: composition and configuration (Cushman et al. 2010; Forman & Godron 1986). The interpretation of a satellite image is being processed using the element of tone, texture, pattern, shape, size, shadow, site and association to derive information about the land cover. This method successfully provides efficient information on temporal trends and spatial distribution of urban areas, which is required for understanding, modelling and projecting land changes (Lu D &Weng Q 2006). The hyperspectral and multispectral images can provide information of inaccessible areas, vegetation patches, related to phenological types, gregarious formations and communities occurring in a unique environmental setup whereas the temporal images help in monitoring a landscape over time (Delcourt 1987; Olorunfemi 1983). The images also provide a digital mosaic of the spatial arrangement of land cover and vegetation types amenable to computer processing (Chuvieco E 1999). The wide variety of land cover changes that occur on the landscape between certain periods of time can be monitored by different change detection methods.

In many landscapes, recent synergistic combinations of natural disturbance and human-related changes have caused disruption and consequently transform the prevailing landforms (Schlesinger et al. 1990; D’Antonio & Vitousek 1992). Therefore, the analyses of the landscape characteristics relative to type and frequency of disturbance can provide implications for land and wildlife management (Harris 1984; Shugart & Seagle 1985; Forman and Godron 1986; Merriam 1988; Turner and Gardner 1991; Callaway and Davis 1993; Turner et al. 1995). The wide variety of land cover changes that occur on the landscape between certain periods of time can be examined by different change detection methods. A wide range of methods is obtainable in remote sensing to analyze these changes, emphasizing different aspects of landscape studies such as land cover conversion, change in vegetation growth, change in the landscape configuration and composition, etc. (Boriana D 2007; Boriana P and Rogan J 2006).

In this study of assessing the landscape structure, principal component analysis of MODIS images was followed by the hybrid classification approach involving both the supervised and unsupervised classification methods and Markov analysis. The accuracy assessment of the classification was accomplished using kappa statistics (Gómez D & Montero J 2011). MODIS (Moderate Resolution Imaging Spectroradiometer) is a payload scientific instrument which collects data in 250m resolution in a 16-day composite. It collects data in 36 spectral bands and because of the 250m resolution, the data covers a large area in a single image (modis.gsfc.nasa.gov). Principal Components Analysis (PCA) invented in 1901 by Karl Pearson and mostly used as a tool in exploratory data analysis to find the most important variables (a combination of them) that explain most of the variance in the data. So, when there is lots of data to be analyzed, PCA can make the task a lot easier. PCA also helps to construct predictive models (Anh T & Magi S 2009). A hybrid classification method was implemented in the analysis. Use of both supervised and unsupervised classification method is ordinarily termed as a hybrid classification (Omo-Irabor O.O.; Oduyemi K 2007). The purpose for using the combination was the large size of the landscape which consists of many landforms. Single classification methods sometimes merge the similar reflectance pixels and thus misclassify the landform. The hybrid classification method prevents this error. Accuracy assessment is a must do the step before implementing any classification methods to know the sources of errors (Powell et al. 2004, Congalton R. G. and Plourde L 2002; Congalton and Green 1999, 1993). The parcel of land can only be in one state at a given time moving successively from one state to the other with a probability which depends only on the current state and not the previous states (Wijanarto A. B. 2006). Analyzed using Markov chain analysis, the probability of moving from one state to the other is called a transition probability which is captured in a transition probability matrix whose elements are non-negative and the row elements sum up to 1 (Camacho O.M.T. et al. 2015; Yang X et al. 2012 and Arsanjani J.J 2011).

This study was carried out in the Central Indian Highland in the state of Gujrat, Rajasthan and Madhya Pradesh. The landscape is a significant tiger habitat having 11 tiger reserves interconnected with each other through corridors (Qureshi et. al 2014). There are 5 established corridors within the landscape (Ranthambore-Kuno-Madhav National Park, Bandhavgarh-Achanakmar, Bandhavgarh-Sanjay-Dubri-Guru Ghasi Das, Kanha-Achanakmar, Kanha-Pench). The landscape supports ∼40% of the total tiger population (Jhala et al. 2011). The result showed the presence of 257 (213-301) tigers in Madhya Pradesh covering 13, 333 sq. km of tiger habitat and 36 (35-37) individuals covering 637 sq. km area in Rajasthan (National Tiger Conservation Authority, 2011). As the area falls under two biogeographic provinces and the river beds of Narmada, this landscape is rich in agricultural productions as well (Eaton R. M 2005; Hugh C 1911). Incidentally, the landscape is the second largest belt of minerals in the country (Prakash, A & Gupta, R. P., 2010) where non-ferrous minerals, uranium, mica, beryllium, aquamarine, petroleum, gypsum and emerald are present in Rajasthan and Gujarat (Vagholikar et al. 2003; Ghose 2003; Swer and Singh, 2004). This also triggers the mining interests (Narain et al. 2005) and falls in the core industrial development zone (Tripathi J.G., 2017). The total population of Rajasthan, Madhya Pradesh and Gujarat as per census 2001 was 5, 65, 07,188; 6, 03, 48,023; 5, 06, 71, 017 with a decadal growth rate 28.4%, 24.3% and 22.7% respectively (Census of India, 2011). In 2011 the total population of these three states gone up to 6,86,21,012; 7, 25,97,565; 6,03,83,628; with a decadal growth rate 21.45, 20.3% and 19.2% (Census India 2011). All the three states have population decadal growth more than the average of India’s decadal population growth in a decade. The area is home to largest scheduled tribe population and among the poorest in the country. High population density, increased need for livelihood facility and economic demand mostly from the mining sector changed the land use and land cover (LULC) of the area (Malaviya, S et al., 2010). The study covering 19 tiger reserves in the landscape showed how the development of urbanization and agricultural activity continued to shrink the tiger habitat over several years (Banerjee 2017). Yadav et al; 2012 examined the condition of Nawegaon-Nagzira corridor where the shrinkage of overall forest cover and water bodies was reported, and the result showed that there has been a decrease in forest and an increase in urban and agriculture. With the agricultural intensification, the constant reduction in forest cover will impact the movement of tigers in this landscape. The outcome of these studies indicated the importance of this landscape in term of wild species conservation and management.

The spatial and temporal change in the land use/cover of this landscape was not assessed before. In this study, we analyzed two different bioprovinces, with different vegetation types and physical parameters. The study focused on two objectives: 1) to assess the land use pattern, and 2) to assess the land use/cover dynamics over time.

### Study Area

The present study includes the Semi-Arid and Deccan Peninsula landscape of India (23°41’9.99”N, 68°24’7.30”E in extreme west to 24°45’41.31”N, 84° 3’25.75”E in extreme east). It falls under two biogeographic provinces of India named as 6A deccan peninsula-central highlands and 4B semi-arid-Gujarat rajputana (Rodgers & Panwar 1988). The area is a part of the longstanding landmass of Northern plains of India and made up of crystalline, Igneous and Metamorphic rocks. The Central Highlands are bounded by Aravalli range on the northwest, Ganga plains on the North & Vindhya Range and Narmada River on the South. It comprises the major portion of the Malwa plateau and is made up of shallow valleys and rounded hills (Krishnan, 1968; Bharucha, 1955; Pramanik et al. 1952 and Thornthwaite, 1948). Certain parts of the landscape are very dry prior to the monsoons when reduced water availability is an issue here for wildlife as well as for people. According to the Champion and Seth (1968) classification, most of this landscape has tropical dry deciduous forests with small sections of tropical moist deciduous forest and tropical thorn forest. The Narmada is considered as a natural boundary between the teak forests of the southern peninsula and sal forests of northern plains (Forsyth 1919). Other than Narmada, Betwa, Parbati, Chambal, Tapi, Son is the other rivers in this landscape. The area includes the state of Madhya Pradesh, Gujarat, and Rajasthan, which consists of 117 districts and 97 cities (Census of India, 2011). Within this zone, several hill ranges are present ranging between elevations of 200m and 1000m such as the Aravalli, Vindhyas where Aravalli is described as the oldest mountain in the world (Walter D. M. et al. 2005). Tropical dry deciduous forests with small sections of tropical moist deciduous forest in the eastern region and tropical thorn forest are the main vegetation type of this landscape. *Syzigium cumini, Murraya paniculata, Sterculia villosa, Tectona grandis, Anogeissus latifolia, Pterocarpous marsupium, Woodfordia fruticosa, Wrightia tictoria, Bauhinia spp, Terminalia alata, Bombax ceiba, Madhuca longifolia, Mitragyna perviflora, Lagerstoemia spp, Accacia pendula, Boswellia sp, Choloroxylon sweitenia* are the common species found in this landscape. As it’s mentioned earlier, the landscape has conservation value as an important habitat for tigers (Panthera tigris).

**Figure 1:**
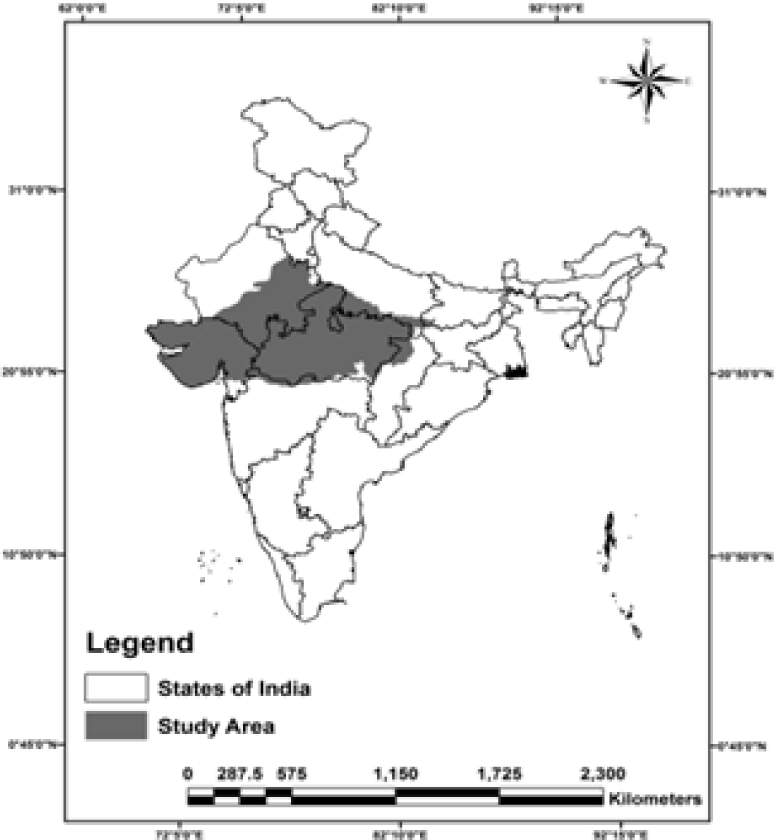
Map of the study area.

## Methodology

MODIS images with 250 m resolution were downloaded (http://earthexplorer.usgs.gov/) for each month of 3 study years (2000, 2006 and 2013). The downloaded images were re-projected from Sinusoidal to the WGS 1984 projection and exported as tiff images by using Arc.GIS 10 (desktop.arcgis.com). Principle component analysis was carried out to convert year wise 12 images to a single image for each year. A hybrid classification using both supervised and unsupervised classification of the images was executed using a signature file, google earth data and information generated from field sampling points using ERDAS Imagine 13 (www.hexagongeospatial.com). The ground truth points were taken from the “All India Tiger Monitoring Project”, 2011 (Jhala et al. 2011). In a total of 448,005 plots were laid covering 478,016 sq. km of forest for collecting information on forest cover (4.1% of the total forested area in India) (Jhala et al. 2011). For other land use classes,’ respectively 200 signature files were prepared using Google Earth. The three principal component images were initially classified into 100 classes in unsigned 8 bit. These classes were then merged into 9 prevalent land use/cover classes in the landscape using a supervised classification approach. This classification approach is coined as a hybrid classification approach which produced 9 Land use/cover classes. The land use classes are as follows:

1. Fallow: represents unused cropland or maybe land used for seasonal crops or land under irrigation.
2. Dense forest: mainly includes moderate to dense canopy forest with 70% or more canopy.
3. Water: represents ponds, rivers and dams.
4. Barren land: representing soil that has almost no vegetation.
5. Scrub: an area of land that is uncultivated and covered with high sparse stunted vegetation.
6. Cropland: percentage of land usually covered with crops.
7. Open Forest: areas adjacent to dense forest, plantation roadside or agricultural field side with 10-30% canopy cover.
8. Desert: a waterless, desolate area of land with little or no vegetation, typically one covered with sand.
9. Settlement: Human habitation.

Accuracy assessment was done for 2000, 2006 and 2013 land cover maps using topographical maps and 870 systematic sampling points.

To achieve the overall objective of painting the present scenario and its progression in the past, present, and near-future, markov chain analysis technique was applied with NDVI classified images. Markov module analyses land cover images and produces a transition probability matrix, a set of conditional probability images. The conditional probability images report the probability that each land cover type would be found at each pixel after the specified number of time units (Lausch A et al. 2015). This was primarily to quantify the changes rather than to spatially locate where the changes are occurring. The change detection used the simple overlay procedure between the classified images of the year 2000 – 2006 and 2013. For this case, an area subdivided into several cells each of which can be occupied by a given type of land use (forest wet/grassland or bare land) at a given time, the transition probabilities were computed on the basis of classification data between time periods which show the probability that a cell would change from one land use type to another within the same particular period in the future (Vázquez-Quintero G. et al. 2016). The transition probability matrix which records the probability of each land cover category to change into the other category was produced by multiplication of each column in the transition probability matrix. In the output of the transition probability matrix, the rows represent the older land cover categories and the column represents the newer categories.

## Results

During the past few decades (2000-2013), the study area has witnessed a substantial increase in built-up areas; it rose from 31317 sq. km. in 2000 to 52338 sq. km in 2013. The increase of area under this class is related to increased rate of economic growth, and industrialization in this landscape.

The fallow land increased from 247645.18 sq. km to 329665.18 sq. km in 2000-2006 but decreased to 226925.56 sq. km in 2006-2013. In the same time frame, the increase in area under settlement (from 31317.81 sq. km in 2000 to 52338.81 sq. km 2013) and barren land (17496.06 sq. km in 2000 to 97969.13 sq. km in 2013) indicates a large amount of agriculture land might have converted into settlements and other development activities in the same time frame. This can also be possible if people have abandoned agriculture at the same time as industrialization was rapidly growing in that landscape (Yadav P.K. et. al. 2012). The similar can be interpreted from the Markov chain analysis result of the time frame 2006-2013 as the probability matrix indicated 18% probability of fallow land changing into barren land. The area of barren land has increased from 17496.06 sq. km to 97969.12 sq. km in 2013. The land under desert class remained consistent throughout the period but the reflectance from perennial water sources and the seasonal farming practices got captured in the classification when few parts of the area got misclassified as fallow or land under water. The cross verification of temporal high-resolution satellite images used in Google Earth (www.google.com/earth) confirmed the same assessment.

The area under dense forest constantly declined in the studied time. Dense forest deteriorated from 84027 sq. km to 79055.56 sq. km (2000 to 2006) and in 2013 the area estimated was 71225.37 sq. km. The probability matrix of dense forest getting converted into agricultural land was 29% in 2000-2006 and 33% in 2006-2013. Open forest occupied an area of 83298.81 sq. km. in 2000, which lessened to 64799.37 sq. km. in 2013. The Markov analysis estimated a probability of changing of open forest types to agricultural land was 46% in 2000-2006 and 32 % in 2006-2013. Cross verification of the classified maps in the time frame of 2000-2013 revealed that dense forest areas present within the tiger reserves also shrank considerably in the landscape. The difference in area is mentioned within the bracket against each tiger reserves such as Melghat (1429.31 sq. km – 1410.68 sq. km), Satpura (1550.81 sq. km. -1222.93 sq. km. from 2000-2013), Pench in Madhya Pradesh part (473.12 sq. km. 450.18 sq. km.), Kanha (1242.31 sq. km. -1208.37 sq. km.), Mukundara Hills (140.37 sq. km. 138.12 sq. km.). Open forest areas in Bandhavgarh (829 sq. km -258.43 sq. km.), Sanjay-Dubri (506.56 sq. km. – 420.31 sq. km) and Panna (795.87 sq. km 387.93 sq. km.). Reduction of dense forest area happened in Ranthambore (236.87 sq. km. – 173.12 sq. km.) from 2006-2013 and change in both the dense (320.56 sq. km. – 231.81 sq. km.) and open forest (506.56 sq. km. 424.87 sq.km) in Sariska was also observed. An urban explosion at the periphery of Mukundara Hills (0.62 sq. km. – 9.56 sq. km) and Ranthambore (12.62 sq. km 19.18 sq. km) was also evident in the time frame of 2006-2013. The scrubland has largely declined from 35713.06 sq. km in 2000 to 23003.88 sq.km in 2013; it might have resulted from the transition of scrubland to the fallow land which was predicted as more than 70% in 2000-2006. The area of scrubland dropped from 2000-2013 in Satpura (6 sq. km-1.37 sq. km), Ranthambore (260.87 sq. km-175.62 sq. km), Mukundara Hills (70.43 sq.km-44.93 sq. km) and in Panna Tiger Reserve (207.93 sq. km 100.12 sq. km) Kappa statistics used to measure classification accuracy was found to be 82.8% (Table 1), showing a significant amount of agreement between the actual and classified land use/cover map.

**Table 1:** Area under different land use/cover classes.

| Class Number | Class Name | The area in sq. km |  |  |
| --- | --- | --- | --- | --- |
|  |  | 2000 | 2006 | 2013 |
| 1 | Fallow | 247645.18 | 329665.18 | 226925.56 |
| 2 | Dense-forest | 84027 | 79055.56 | 71225.37 |
| 3 | Water | 14185 | 16170.75 | 6360.75 |
| 4 | Barren- land | 17496.06 | 19664.43 | 97969.12 |
| 5 | Scrub | 35713.06 | 18174.12 | 23003.87 |
| 6 | Crop | 159399.43 | 129854.62 | 132713.37 |
| 7 | Open-forest | 83298.81 | 59472.93 | 64799.37 |
| 8 | Settlement | 31317.81 | 22341.31 | 52338.81 |

**Table 2:** Kappa statistics showing the accuracy of the image classification.

| ACCURACY TOTALS |  |  |  |  |  |
| --- | --- | --- | --- | --- | --- |
| Class | Reference | Classified | Number | Producers | Users |
| Name | Totals | Totals | Correct | Accuracy | Accuracy |
| Fallow | 7 | 35 | 6 | 85.71% | 17.14% |
| Dense Forest | 78 | 58 | 58 | 74.36% | 100.00% |
| Water | 17 | 15 | 15 | 88.24% | 100.00% |
| Barren land | 74 | 62 | 57 | 77.03% | 91.94% |
| Scrub land | 37 | 39 | 22 | 59.46% | 56.41% |
| Crop land | 6 | 10 | 5 | 83.33% | 50.00% |
| Open forest | 91 | 92 | 78 | 85.71% | 84.78% |
| Settlement | 97 | 96 | 96 | 98.97% | 100.00% |
| Overall Classification Accuracy = 82.80% |  |  |  |  |  |

Identification of respective land use/cover classes and the accuracy of the classified images of 2000 and 2006 were assessed by identifying bigger patches (more than 1ha) and using ground-truth points of 2013, distributed in those bigger patches.

### Transition Probability Matrix

As seen in Table 3 the fallow land had a 63% probability of remaining in the same class while there was a chance of 24% of the fallow land getting converted into cropland. Similarly, the dense forest had a 19% probability of changing into fallow land and 10% probability of getting converted into cropland whereas there was a 19% probability of getting converted into an open forest. Summing up, in this period the dense forest had a probability of 29% of getting converted into agricultural land use. It is predicted that a considerable amount of scrubland could also be converted into the fallow land (71%) in the same time frame. In the later phase a considerable amount of scrub (33%) was predicted to be converted into fallow land while 12% of the dense forest was predicted to be converted to fallow with another 23% could get converted into open forest category.

**Table 3:** Transitional Probability table derived from the land use land cover map of 2000-2006.

| Classes | Fallow | Dense-forest | Water | Barren-land | Scrub | Crop | Open forest | Settlement |
| --- | --- | --- | --- | --- | --- | --- | --- | --- |
| Fallow | 0.63 | 0.04 | 0.00 | 0.01 | 0.03 | 0.24 | 0.04 | 0.01 |
| Dense-forest | 0.19 | 0.49 | 0.00 | 0.00 | 0.01 | 0.10 | 0.19 | 0.00 |
| Water | 0.04 | 0.00 | 0.75 | 0.18 | 0.00 | 0.00 | 0.01 | 0.00 |
| Barren-land |  | 0.01 | 0.18 | 0.60 | 0.00 | 0.00 | 0.02 | 0.00 |
| Scrub | 0.71 | 0.04 | 0.00 | 0.04 | 0.06 | 0.06 | 0.08 | 0.01 |
| Crop | 0.58 | 0.05 | 0.00 | 0.00 | 0.03 | 0.29 | 0.04 | 0.00 |
| Open forest | 0.36 | 0.23 | 0.00 | 0.00 | 0.05 | 0.10 | 0.25 | 0.01 |
| Settlement | 0.18 | 0.03 | 0.01 | 0.00 | 0.00 | 0.15 | 0.03 | 0.60 |

**Table 4:** Transitional Probability table derived from the land use land cover map of 2006-2013.

|  | Fallow | Dense-forest | Water | Barren-land | Scrub | Crop | Open-forest | Settlement |
| --- | --- | --- | --- | --- | --- | --- | --- | --- |
| Fallow | 0.43 | 0.03 | 0.00 | 0.18 | 0.05 | 0.20 | 0.06 | 0.06 |
| Dense-forest | 0.12 | 0.47 | 0.00 | 0.03 | 0.01 | 0.11 | 0.23 | 0.01 |
| Water | 0.14 | 0.01 | 0.19 | 0.61 | 0.00 | 0.00 | 0.02 | 0.02 |
| Barren-land | 0.19 | 0.00 | 0.05 | 0.72 | 0.01 | 0.00 | 0.02 | 0.01 |
| Scrub | 0.33 | 0.03 | 0.00 | 0.18 | 0.14 | 0.15 | 0.17 | 0.00 |
| Crop | 0.39 | 0.06 | 0.00 | 0.04 | 0.01 | 0.36 | 0.05 | 0.07 |
| Open-forest | 0.20 | 0.27 | 0.00 | 0.07 | 0.04 | 0.12 | 0.27 | 0.02 |
| Settlement | 0.01 | 0.01 | 0.00 | 0.00 | 0.01 | 0.01 | 0.04 | 0.92 |

**Figure 2:**
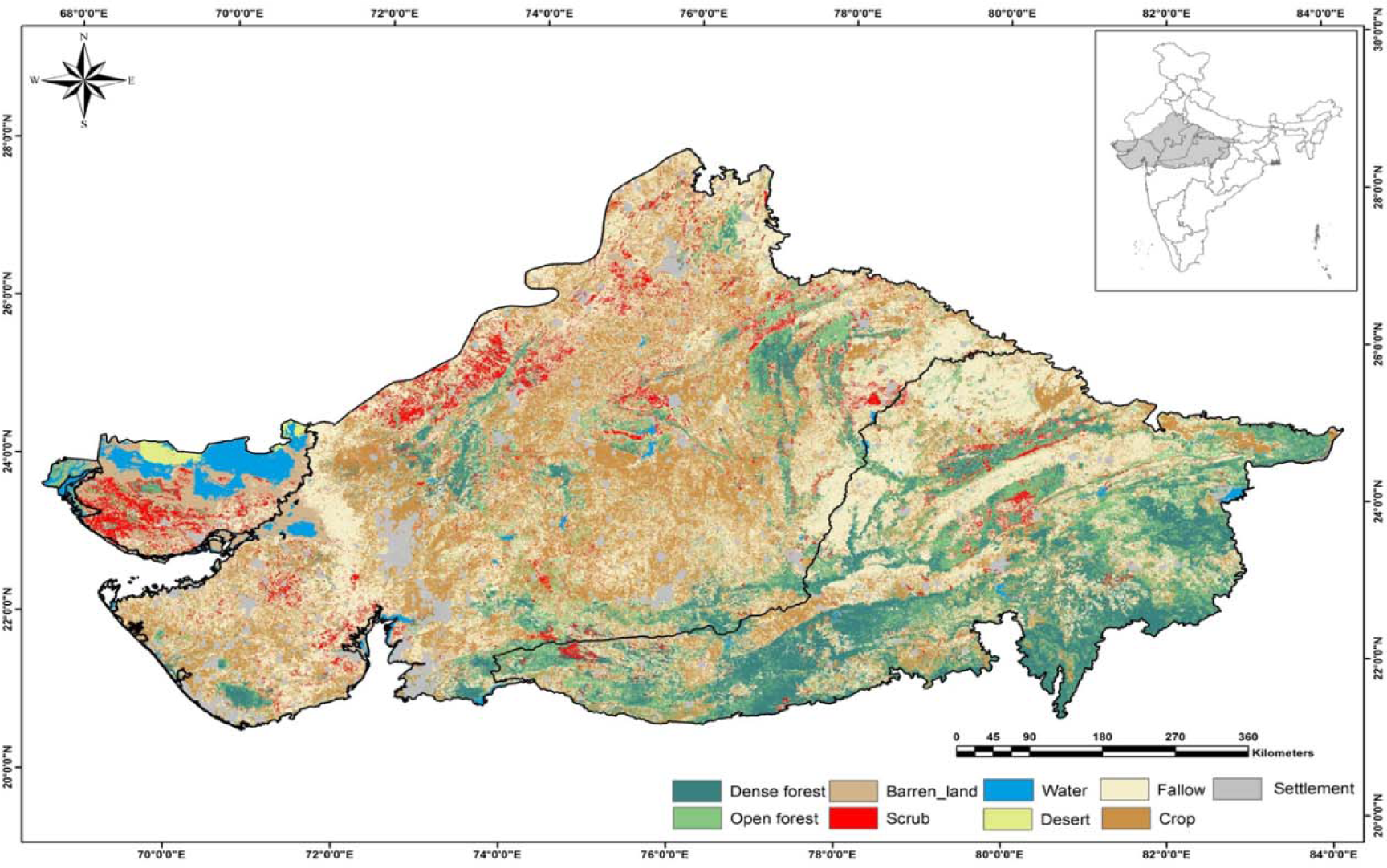
Land use land cover map of the year 2000.

**Figure 3:**
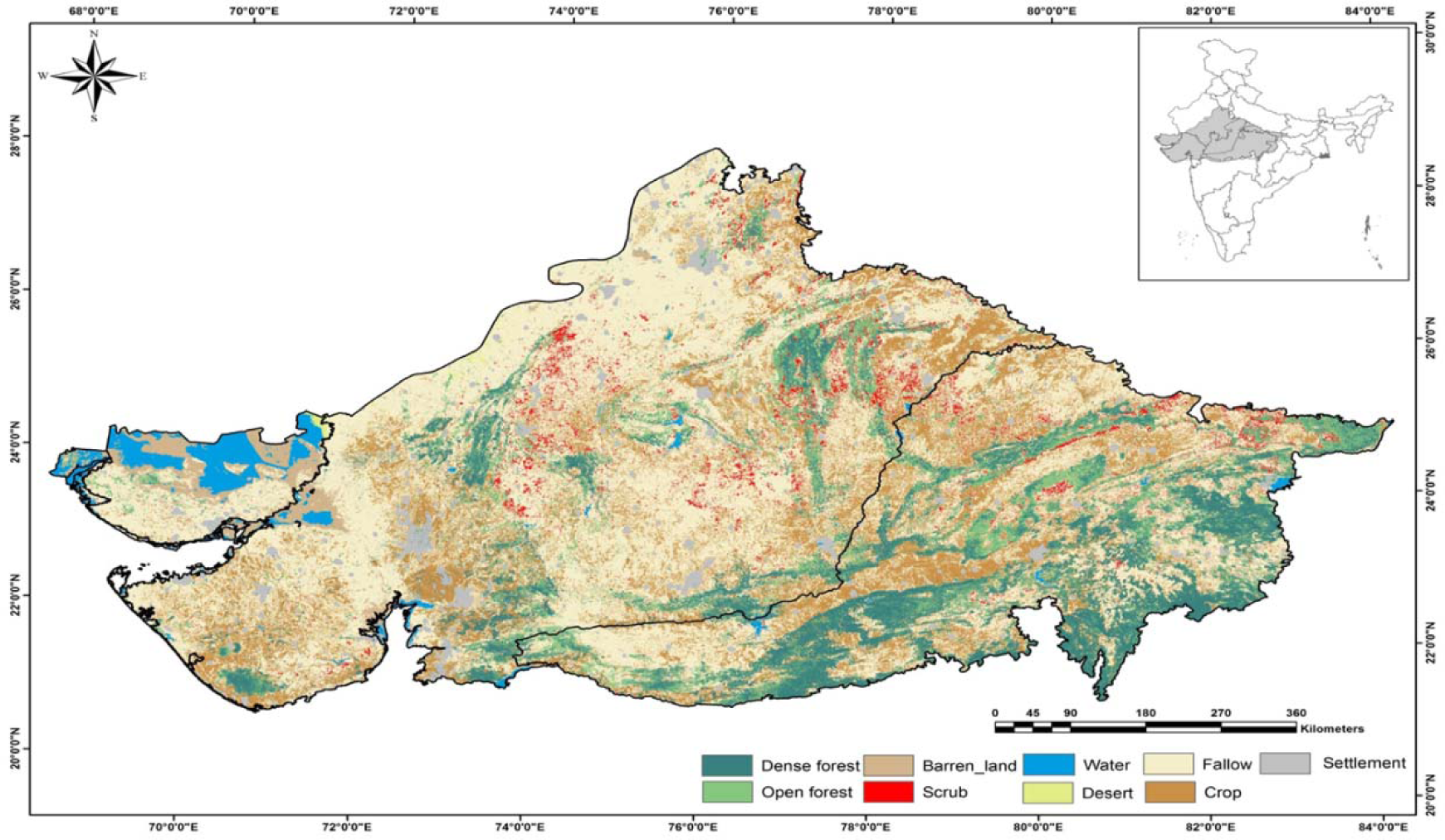
Land use land cover map of the year 2006.

**Figure 4:**
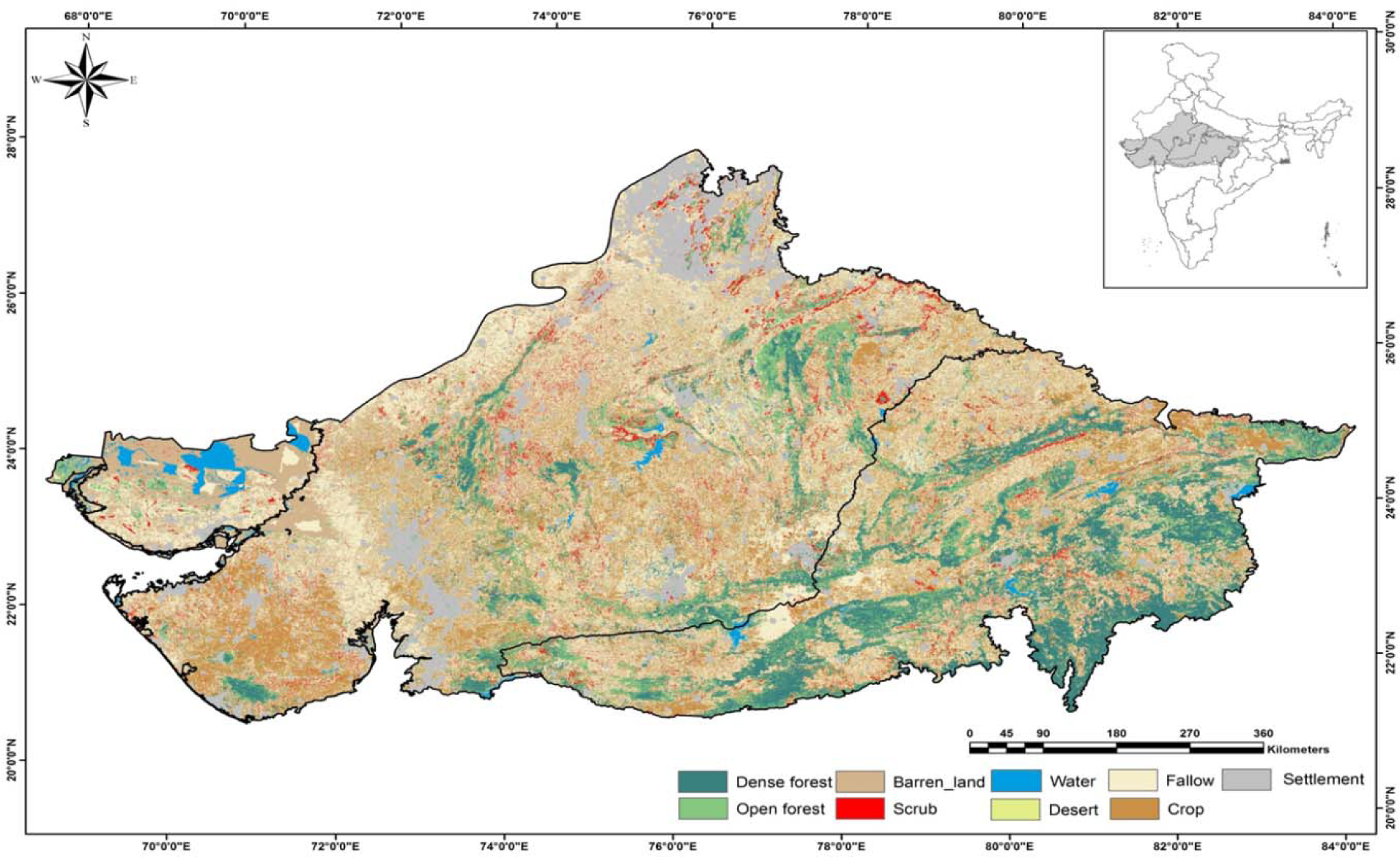
Land use land cover map of the year 2013.

## Discussion

The total population of Rajasthan, Madhya Pradesh and Gujarat showed an increasing trend as per Census data and the decadal growth rate was also high from the average decadal growth of the country. The final LULC map of 2013 showed the northern and southwestern part of the landscape witnessed speedy growth in the settlement areas which resulted in the conversion of agricultural land, fallow land and scrubland into such type of land class. The economic development within the area due to industrialization may have attracted colonization of labours in seek of a better livelihood. That might have been the reason behind the rise in settlements in this area to provide shelter to this migrant population. Expansion of the buildup areas near to the tiger reserves is mentioned in the results earlier. Removal of open forest for the residential areas is less time consuming and availability of the woods for the household works also gets fulfilled. Therefore, it is one of the causes of declination of open forest areas. Also, paper production industries are present in the landscape. These companies try to establish workshops near to the forest areas to get softwood easily and to cut down the transport cost. An open forest area is effortlessly accessible and thus causes the encroachment rapidly.

The industrialization in the area provided an opportunity of improving the economic status of livelihood of the native people and low agricultural productivity could be the reason behind the per cent increase in barren land. All these have led the way to large-scale mining industries in this area. Malaviya S; et al. 2009 found a significant decline in forest cover, especially of the Sal-mixed forests. Mining caused a lot of environmental pollution due to its continuous process of the blast and digging. Indian Bureau of Mines (IBM) had identified abandoned/orphaned mines which had been left un-reclaimed prior to the promulgation of rules about Mine Closure Plan in April 2003. Through a special study at the national level, 297 abandoned mine sites were identified of which 106 abandoned mine sites belonged to Public Sector Undertakings. Among the 106 sites, 24 mine sites became operational again. Thus 82 abandoned sites require reclamation and rehabilitation now. Of this 82, 28 sites were present in Madhya Pradesh, 3 in Rajasthan and 1 in Gujarat (ibm.gov.in). As it is evident, in 2000-2006-time frame, most of the agricultural land remained fallow (63%) whereas the probability of crop land to fallow was 0.58 which means more than 50% of the crop land also remained uncultivated during this time frame. As the mining industries gave the opportunity to thousands of people to work as daily wages and it might have more profitable to them than the agriculture also can be one the cause behind less cultivates in the region. Added to this, more than 70% of the scrubland showed probability of getting converted into the fallow land. This altogether gave rise to the amount of area under fallow land in 2000-2006 timeframe. To its contrary, in 2006-2013-time frame, though the land under crop increased reducing the area under fallow, a considerable amount of land under fallow showed tendency to be converted in under crop (0.20). Added to this 18% of the fallow land further degraded to barren land and settlement (6%). This altogether has reduced the area under cultivation (fallow) in 2006-2013-time frame under influence of urban explosion as well as cropping.

As the area has potentiality and resources to support large mammals it needs more assessments to conserve the wild population. The disturbance for a longer time can affect the present ecosystem dynamics (Sousa 1984) and species diversity (Connell 1978; Miller 1982; Turner 1987; Caswel and Cohen 1993), coexistence (Wiens 1977; Palmer 1992). With such a prominence in the big cat conservation, the landscape needs a more concentrated study to protect the vulnerable ecosystem. Environmental planning with a landscape approach has potential to generate essential information for understanding ecological dynamics and to consolidate management proposals for landscapes with remnants of natural vegetation in areas with ongoing human activities (Metzger; Brancalion, 2013).

## Conclusion

The study conducted in the Deccan Peninsula and Semi-Arid bioprovinces advocates that multi-temporal satellite imagery plays a fundamental role in quantifying spatial and temporal phenomena which are else not possible to attempt through conventional mapping. The remote sensing and GIS tools have been applied to analyze land use/land cover changes in this landscape. Nine major LULC classes have been clearly identified from the area. The study also provided the trend of foremost changes in the land use/land cover classes of the study area during the time frame 2000 to 2013. It is observed that there is an impression of industrialization, the development of urban areas and agricultural commotion on forest cover along with other factors. Shrinkage of forest near and within the protected areas will have a negative impact on the ecosystem and wildlife species consisted there. As mentioned in the results, expansion of the residential areas near to the protected areas caused a lot of disruption. This penetration in a consistent frequency in the habitats of the wild animals often divert animals from their original dispersal route and enter nearby suburban areas leading to conflict situations. The present study illustrates the impact of the different anthropogenic activity on land use/cover change in a large landscape. These changes can initiate an exertion to the wildlife conservation. This study can facilitate generating the base level information of the areas to focus more from the conservation point of view to the state forest departments and lawmakers. The more intense study can be done to quantify the major changes in the specific protected areas. Settlements near to the buffer areas of the tiger reserves need to be restricted to prevent encroachment and that can protect the forest cover in the buffer areas. Therefore, a landscape level sustainable framework of management intervention can improve the habitat quality of the existing protected areas in this landscape.

